# Cryo-ET of IgG bivalent binding on SARS-CoV-2 provides structural basis for antibody avidity

**DOI:** 10.1101/2025.02.28.640788

**Authors:** Hangping Yao, Yutong Song, Qi Huang, Miaojin Zhu, Jiaming Liang, Zheyuan Zhang, Xiaodi Zhang, Dongyang Dong, Danrong Shi, Zhigang Wu, Xiangyun Lu, Haibo Wu, Yong Chen, Sai Li

## Abstract

The bivalent nature of IgG enhances its neutralization potency against enveloped viruses; however, on-virion structural details of IgG bivalent binding with antigens remain elusive. Here we used cryo-ET to investigate how two potent IgGs P17 and S309 interact with S-trimers on the SARS-CoV-2 surface. We found that these antibodies exploit the mobility of S-trimers to form diverse bivalent binding patterns. P17 stabilizes S-trimers in a one-RBD-up conformation and gathers S-trimer into linear multimers within minutes, whereas S309 primarily forms circular S-trimer assemblies that extend into lattice-like structures. Additionally, both IgGs can facilitate inter-virion coupling through bivalent binding of opposing S-trimers. These findings provide a structural basis for understanding IgG avidity and offer insights for antibody engineering and vaccine design.

## Introduction

IgG, the most abundant antibody isotype in serum, plays a crucial role in the humoral immune response against a wide array of pathogens, including enveloped viruses. As a key component of adaptive immunity, IgG contains two antigen-binding fragments (Fab regions)^1^, enabling simultaneous engagement with two epitopes. This bivalent binding enhances overall binding strength, a phenomenon termed avidity. Avidity improves viral neutralization efficiency by enhancing epitope recognition, promoting multivalent interactions, restricting viral escape, and facilitating immune clearance. Its role in viral neutralization has been reported across various enveloped viruses, including influenza virus^2^, respiratory syncytial virus (RSV)^3^, MERS-CoV^4^, Chikungunya virus^5^ and Dengue virus^6^. The avidity effects on SARS-CoV-2 are particularly evident when comparing the neutralization efficiency of some potent IgGs to their Fabs fragments^7^, as well as the overall potency of bivalent IgGs to monovalent IgGs^8^. Notably, certain IgGs induce the shedding of S1 subunits, whereas their Fab fragments do not^9^. In addition to the Fab region, IgG avidity is also conferred by the Fc region^10^. This region interacts with Fc receptors (FcRs) on immune cells, facilitating complement activation, antibody-dependent cellular cytotoxicity (ADCC), and phagocytosis, thereby enhancing viral clearance^11^. However, interactions between a single Fc region and FcRs are relatively weak, necessitating IgG multimerization into immune complexes to achieve sufficient avidity, strengthen FcR engagement, and effectively trigger effector functions^12^.

Despite extensive functional research, the structural basis of high-avidity IgG-antigen interactions on enveloped viruses remains largely unresolved. This challenge stems from the inherent flexibility of IgG molecules, originating from their hinge regions^13^, and the pleomorphic nature of enveloped viruses^14,15^. Previous studies employing X-ray crystallography and single-particle cryo-electron microscopy (cryo-EM) have sought to overcome these challenges by analyzing Fab complexes with viral glycoprotein (GP) fragments such as peptides^16,17^, receptor binding domains (RBDs)^18,19^, or GP ectodomains^20–23^. While these approaches have yielded critical insights into Fab-epitope interactions, the physiologically relevant context imposed by viral membranes is missing. Investigations utilizing Fab fragments on intact enveloped viruses provide a more native setting; however, these approaches overlook the manner in which IgG molecules engage multiple antigenic sites on the viral surface—an essential aspect of high-avidity binding^24–26^. To date, only a limited number of structural studies have examined intact enveloped viruses incubated with full-length IgGs, and most of these have reported observations restricted to single Fab binding events^27,28^. These findings suggest that IgG bivalency is context-dependent, influenced by IgG structure, epitope accessibility, and the spatial arrangement of antigens on the viral surface.

To address these limitations, we employed cryo-electron tomography (cryo-ET) and subtomogram averaging (STA) to elucidate how two potent IgGs, P17 and S309, engage with S-trimers on SARS-CoV-2 surface. P17 and S309 are class-2 and class-3 anti-RBD antibodies^29^, respectively, both target the RBD in up- or closed-conformations and exhibit high neutralization potency against SARS-CoV-2^30,31^. Our findings demonstrate that the IgG form of P17 (P17-IgG) possesses greater neutralization potency than its Fab fragment (P17-Fab). Likewise, the IgG form of S309 (S309-IgG) achieves complete neutralization, whereas its Fab counterpart (S309-Fab) does not^31^, highlighting the crucial role of bivalency in antiviral efficacy. Cryo-ET and STA analysis revealed diverse inter-spike bivalent binding modes in SARS-CoV-2 virions incubated with P17 or S309. Both IgGs utilize their bivalent nature and the intrinsic mobility of S-trimers to assemble S-IgG dimer-of-trimers complexes. When mapped onto the viral surface, these S-IgG complexes were found actively clustering S-trimers into higher-order S-IgG assemblies. Notably, both IgGs are also capable of inter-virion coupling by bivalently binding opposing S-trimers on neighboring virions. Collectively, these findings establish a structural framework for IgG avidity and provide valuable insights for antibody engineering and vaccine development, particularly in the optimization of epitope presentation and multimerization strategies to enhance immune responses.

## Results

### P17-IgG Binds All RBDs on-Virion and Alters the Conformational Ratio of Prefusion S

To investigate the neutralizing mechanisms of P17-IgG, we first examined its binding to S-trimers on SARS-CoV-2 and its impact on spike conformations. The SARS-CoV-2 WT strain (ID: ZJU-05)^32^ was propagated in a BSL-3 laboratory, where both P17-Fab and P17-IgG exhibited potent neutralization of live SARS-CoV-2 virions. The half-maximal inhibitory concentration (IC_50_) of P17-IgG was 25.2 ng/mL, nearly twice as potent as P17-Fab, which had an IC_50_ of 46.3 ng/mL (Fig. 1A).

**Fig. 1.**
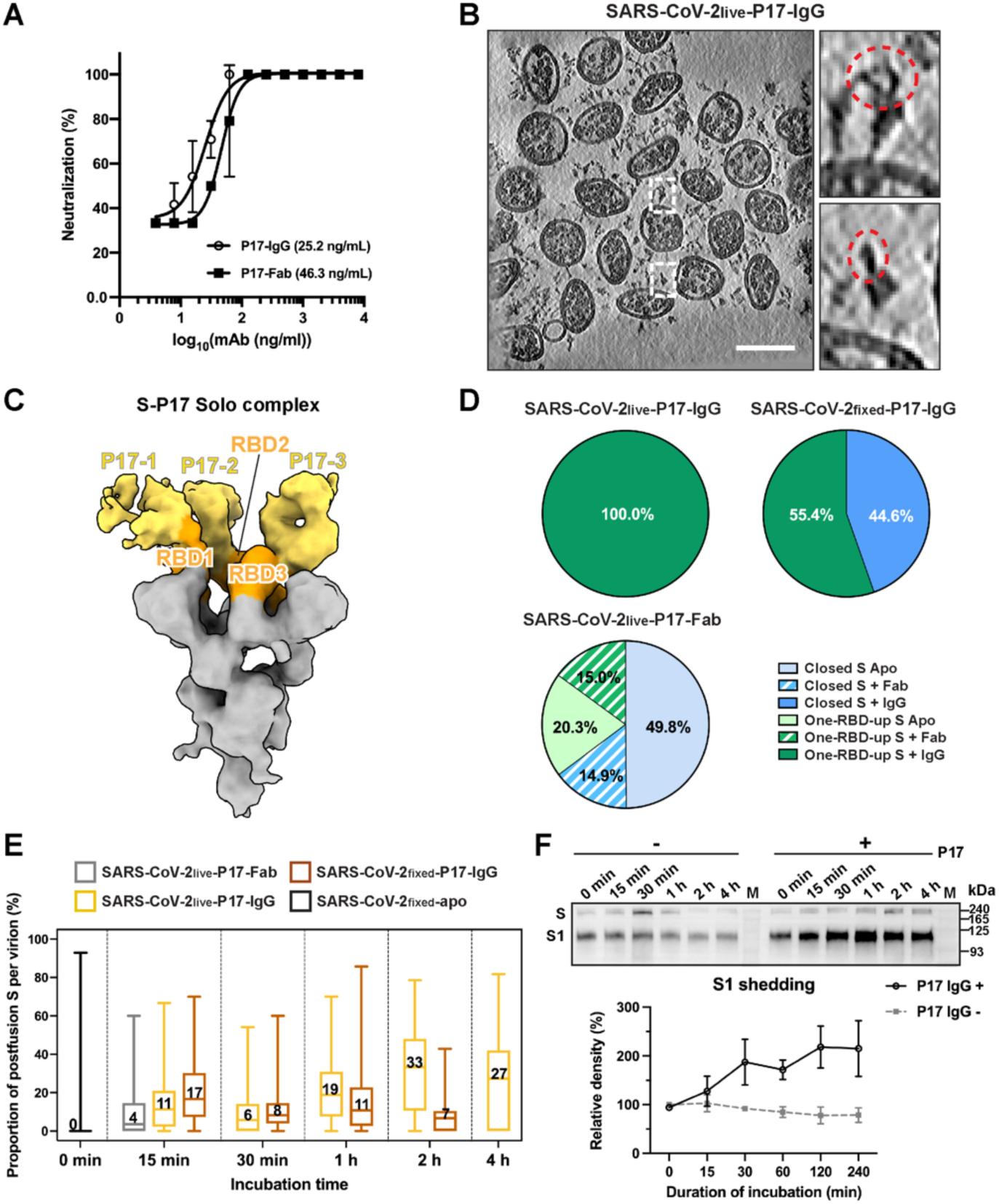
Neutralizing antibody P17 stabilizes SARS-CoV-2 S in one-RBD-up conformation and induces S1 shedding. (**A**) Neutralization efficiency of P17-IgG and P17-Fab against WT SARS-CoV-2 measured as a function of antibody concentration. Data are presented as mean ± s.d. from four independent experiments. (**B**) An exemplary tomogram slice (5 nm thickness) of prefusion S (white boxes) on SARS-CoV-2_live_-P17-IgG virions (15 minutes incubation), showing additional P17 densities (red circles) near the RBDs. Scale bar: 100 nm. (**c**) On-virion structure of the S-P17 Solo complex determined from prefusion S of SARS-CoV-2_live_-P17-IgG across all incubation durations. Each RBD (orange) is bound to a Fab fragment (yellow) of P17-IgG. The up-RBD is numbered RBD1, with its bound P17 numbered P17-1, the remaining RBDs and P17 are numbered sequentially in a clockwise direction. (**D**) Conformation ratios of prefusion S across different samples. SARS-CoV-2_live_-P17-IgG exhibited a predominance of the one-RBD-up conformation with all RBDs bound with P17 (One-RBD-up S + IgG), whereas SARS-CoV-2_fixed_-P17-IgG and SARS-CoV-2_live_-P17-Fab displayed mixed conformations, including closed S bound with IgG (Closed S + IgG), closed S bound with Fab (Closed S + Fab), unbound closed S (Closed S Apo), one-RBD-up S bound with IgG (One-RBD-up S + IgG), one-RBD-up S bound with Fab (One-RBD-up S + Fab) and unbound one-RBD-up S (One-RBD-up S Apo). (**E**) Proportion of postfusion S per virion. SARS-CoV-2_live_-P17-IgG showed a time-dependent increase in the proportion of postfusion S, as compared to SARS-CoV-2_fixed_-P17-IgG. Boxplots showing the median (center line), interquartile range (box), and whiskers indicating the minimum and maximum values.(**F**) Western blot analysis of P17-induced S1 shedding. HEK293T cells expressing S on the surface were incubated with P17-IgG (2 µg/mL) for up to 4 hours. Western blotting of cell supernatants detected S1 fragments using an anti-S1 NTD antibody. Quantification of S1 shedding (bottom) was normalized to the 0-minute P17(-) condition and presented as mean ± s.e.m. from three independent experiments.

To elucidate the structural basis underlying this enhanced neutralization, we employed cryo-ET and STA to analyze SARS-CoV-2 incubated with either P17-IgG or P17-Fab (table S1). The supernatant from infected cells, at a concentration of approximately 10⁷ virions/mL, was incubated with P17-IgG (2 μg/mL) at room temperature for 15 min, 30 min, 1 h, 2 h or 4 h followed by paraformaldehyde (PFA) inactivation (SARS-CoV-2_live_-P17-IgG). To compare the effects of IgG and Fab binding on virions, live virions were incubated with P17-Fab in the same molar Fab concentration (1.33 μg/mL) for 15 min (SARS-CoV-2_live_-P17-Fab) as control. Notably, the ratio of virions to IgG/Fab for the cryo-ET sample preparation is significantly higher than that of the neutralization assay. Since PFA fixation is not an immediate process, experimental observations should be interpreted in relative terms rather than absolute time points. To assess the effect of chemical fixation on the antigenicity of on-virion S-trimers, control samples of concentrated prefixed virions (SARS-CoV-2_fixed_-apo) were incubated with P17-IgG (43.5 µg/mL) for durations of 15 min, 30 min, 1 h or 2 h (SARS-CoV-2_fixed_-P17-IgG). Following inactivation validation, samples were transferred to a BSL-2 lab for purification and imaging.

Using cryo-ET, we imaged and reconstructed 3,258 SARS-CoV-2_live_-P17-IgG virions, 599 SARS-CoV-2_live_-P17-Fab virions and 885 SARS-CoV-2_fixed_-P17-IgG virions as tomographic data. Tomographic analysis revealed that the viral surface exhibited more complex S-trimers architectures than those observed on SARS-CoV-2_fixed_-apo^32^. The densities of S-trimers appear larger, with additional densities protruding from S-trimers’ head regions, indicative of P17 binding (Fig. 1B). Subsequent STA of 29,981 prefusion S on SARS-CoV-2_live_-P17-IgG demonstrated that all prefusion S were engaged with P17 and were exclusively in a one-RBD-up conformation. On the prefusion S structure, each of the three RBDs, regardless of being in the up- or down conformations, is bound with P17. Further classification of the spikes revealed an S-P17 Solo complex structure (Fig. 1C). To determine whether any two of the three Fab densities within an S-P17 Solo complex could originate from a single P17-IgG, previously reported class-2 RBD-neutralizing antibody Fab model (PDB: 7K8O)^29^ into the Fab densities. Given the hinge length constraints of an IgG, the maximal C-terminal Fab CH1 domain (residue 222 of the heavy chain) separation should not exceed 65 Å^29^. However, the distances measured in the S-P17 Solo complex were 81, 100, and 146 Å (fig. S1A), confirming that each Fab density belonged to separate P17-IgG molecules. For convenience of description, the up-RBD is numbered as RBD1 with its attached P17 numbered as P17-1, the remaining RBDs and P17 molecules are numbered ascendingly in the clockwise direction. The S-P17 Solo complex structure was highly consistent with the recombinant structure of S in complex with P17-Fab (PDB: 7CWM) (fig. S2A).

Given that approximately half of the prefusion S-trimers on SARS-CoV-2_fixed_-apo adopt a closed conformation^32^, we hypothesized that P17-IgG binding stabilizes prefusion S in a one-RBD-up conformation. To validate this, we first performed STA on 10,377 prefusion S of SARS-CoV-2_fixed_-P17-IgG. Despite all RBDs being occupied by P17, only 55.4% prefusion S adopted a one-RBD-up conformation, with the remainder adopting a closed conformation (Fig. 1D and fig. S2C). This conformational ratio mirrored that of prefusion S on SARS-CoV-2_fixed_-apo. Next, STA on 12,210 prefusion S from SARS-CoV-2_live_-P17-Fab revealed that over half of the prefusion S remained unbound. Among the bound spikes, which displayed both one-RBD-up and closed conformations, RBD occupancy was incomplete (Fig. 1D and fig. S2B). Compared to recombinant S-P17-Fab structures exhibiting one-RBD-up or two-RBD-up conformations^30^, our on-virion S-P17-Fab Solo complexes on SARS-CoV-2_live_-P17-Fab displayed different conformational distributions and significantly lower P17-Fab binding efficiency. Notably, a construct of prefusion-stabilized ectodomain S that exclusively adopts a one-RBD-up conformation^33^ were used for recombinant studies.

Intriguingly, the presence of P17-IgG correlated with an increased proportion of postfusion S on SARS-CoV-2_live_-P17-IgG. To investigate whether P17-IgG induces S1 shedding, we quantified pre- and postfusion S on SARS-CoV-2_live_-P17-IgG, SARS-CoV-2_live_-P17-Fab and SARS-CoV-2_fixed_-P17-IgG virions. Statistical analysis demonstrated that the proportion of postfusion S over the total number of S per SARS-CoV-2_live_-P17-IgG virion is markedly higher than that of SARS-CoV-2_fixed_-apo. Additionally, this proportion increased over incubation time on SARS-CoV-2_live_-P17-IgG, but remained less changed on SARS-CoV-2_fixed_-P17-IgG (Fig. 1E). Western blot analysis of HEK293T cells expressing S protein confirmed enhanced S1 shedding upon P17-IgG treatment (Fig. 1F), supporting the hypothesis that P17-IgG facilitates S1 dissociation, which may contribute to neutralization (Fig. 1F).

Collectively, these findings demonstrate that P17-IgG efficiently binds all three RBDs within an on-virion S-trimer, irrespective of its conformational state or fixation status, whereas P17-Fab exhibits lower binding efficiency. Upon binding with an unfixed on-virion S-trimer, P17-IgG, but not P17-Fab, stabilizes all prefusion S in a one-RBD-up conformation. By stabilizing the one-RBD-up conformation and inducing S1 shedding, P17-IgG exerts its neutralizing effects through both avid binding and conformational modulation.

### Bivalent Binding of P17-IgGs Forms Eleven Modes of S-P17 Gemini Complexes and Gathers S-Trimers into Strings

When inspecting the averaged structure of all prefusion S on SARS-CoV-2_live_-P17-IgG at lower contour thresholds, extended densities emerged from the RBD1-P17-1 and RBD2-P17-2 domains, forming a second S-P17 complex. Interestingly, the connecting densities between these S-P17 complexes exhibited characteristic feature of two IgG molecules, with two pairs of Fab densities docking across adjacent S-trimers to form a symmetric dimer-of-trimers structure. To further investigate this phenomenon, we re-aligned the complexes with the dual S-P17 complexes centered and applied dimeric symmetry. Subsequent classification identified two distinct configurations: the S-P17 Solo complexes (Fig. 1C) (48.6% prefusion S, 11.4 Å resolution) and the S-P17 Gemini complexes (51.4% prefusion S, 12.4 Å resolution). To validate the presence of S-P17 Gemini complexes on virions, we examined the viral surface in the tomograms and identified pairs of prefusion S bridged by IgG-like densities (Fig. 2A). Within the Gemini complex, two Fabs pairs (Fab1-Fab2 of P17-1 and P17-2 in top view) couple the two S-trimers on their RBD1s and RBD2s, forming two separate bridges. The third RBD of each S-trimer was occupied by a separate P17 (P17-3 and P17-4), although the Fab densities there was weaker than those observed in the RBD1- or RBD2-bound regions. The Fc regions were largely absent from all P17 densities, likely due to their inherent mobility relative to the Fab fragments (Fig. 2B). To determine whether any two neighboring Fab fragments within the Gemini complex originated from a single P17-IgG molecule, we fitted a previously reported class-2 RBD-neutralizing antibody Fab model (PDB: 7K8O)^29^ into the Fab densities (fig. S1B). Two key criteria were used for validation: (1) the distance between the Fab fragments must not exceed 65 Å^29^, as constrained by the IgG hinge region, and (2) the orientations of the two Fabs shall converge on their C-termini. The measured distance between the heavy chain residues 222 of the fitted models was 41 Å, aligning with the analogous distance observed in the intact human IgG1 (PDB: 1HZH)^1^. Together with the continuous density between the Fab fragments and the converging orientations of their C-termini, these findings confirmed that the two S-trimers were coupled by two P17-IgG molecules (P17-1 and P17-2) within the S-P17 Gemini complex.

**Fig. 2.**
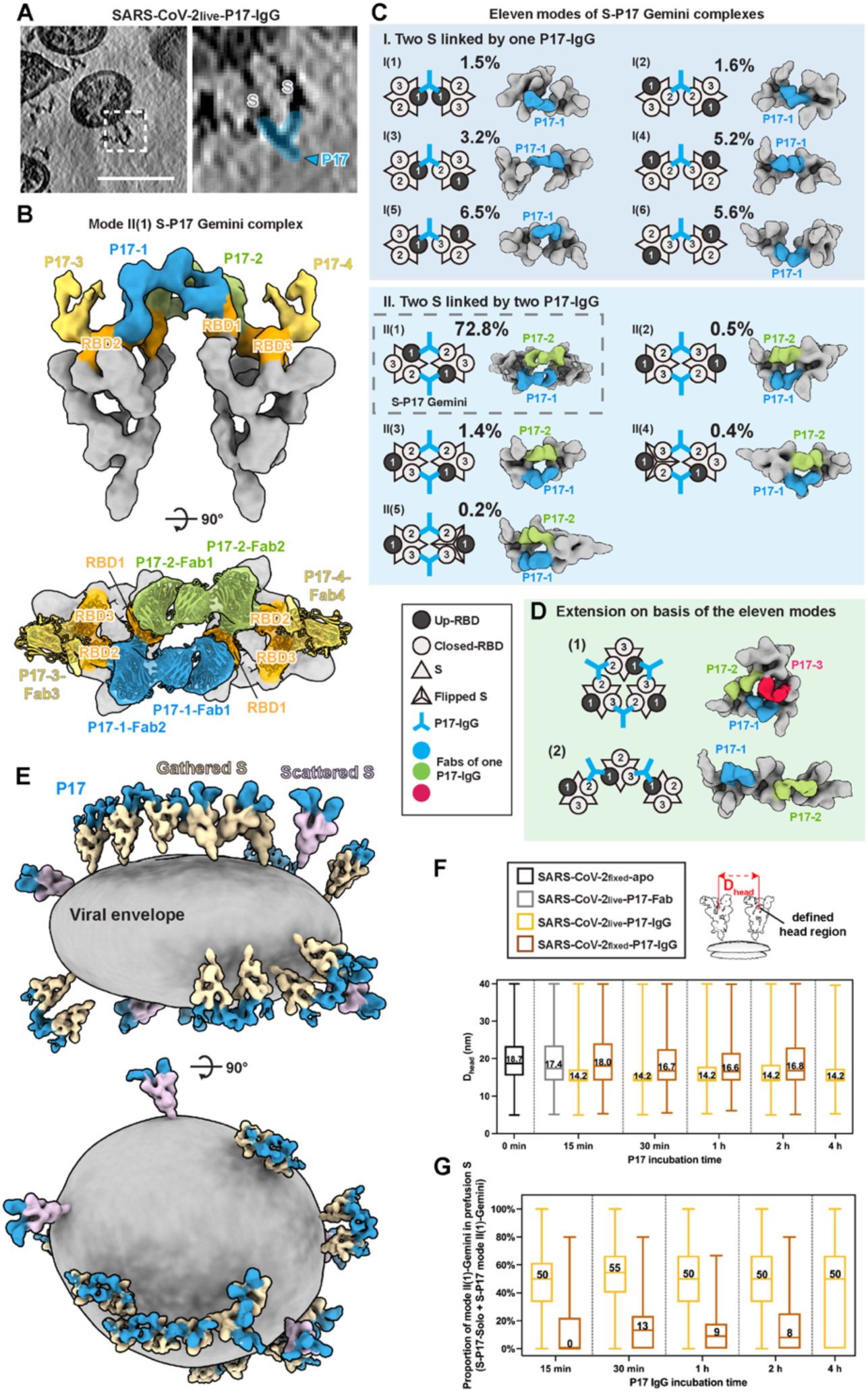
Bivalent binding of P17-IgG forms S-P17 Gemini complexes and gathers S-trimers into strings. (**A**) An exemplary tomogram slice (5 nm thickness) of the SARS-CoV-2_live_-P17-IgG (15 minutes incubation) shows an S-P17 Gemini complex (white box), where two S-trimers are bridged by an IgG (light blue). Scale bar: 100 nm. (**B**) On-virion structure of the S-P17 Gemini complex determined from prefusion S of SARS-CoV-2_live_-P17-IgG across all incubation durations. Two S-trimers (gray, with RBDs in orange) are coupled by two P17-IgG molecules (light blue and green). Each P17-IgG forms bivalent interactions between RBD1 and RBD2 of the adjacent S-trimers. The third RBD of each S-trimer is occupied by a Fab fragment of a separate P17-IgG (yellow). In the top view, the RBDs from PDB: 7CWM and Fab models from PDB: 7K8O were fitted to the density of RBD and Fab in the S-P17 Gemini complex structure. (**C**) Schematics, their exemplary composite structures and proportions of the 11 modes of S-P17 Gemini complexes. These complexes were found by projecting the S-P17 Solo structures onto their refined coordinates on SARS-CoV-2_live_-P17-IgG virions. These modes were grouped into two categories based on the number of coupling P17-IgG molecules: Category I (single IgG bridges two S-trimers) and Category II (two IgGs bridge two S-trimers). Mode II(1) corresponds to the resolved S-P17 Gemini complex in (B). (**D**) Schematics and their exemplary composite structures of the extended S-P17 multimers found on SARS-CoV-2_live_-P17-IgG. The majority of multimers are linearly formed by P17-IgG bivalent binding, only few are in the form of trimer-of-trimers. (**E**) A composite structure of a representative SARS-CoV-2_live_-P17-IgG virion (15 minutes incubation) was reconstructed by mapping all S-P17 complexes onto the viral envelope. S-trimers gathered into clusters are colored wheat, while scattered S-trimers are colored thistle. P17 are colored light blue. The viral envelope (grey) was reconstructed by segmenting manually labeled densities. (**F**) Distance between the head regions of neighboring S-trimers (D_head_) under different conditions. The D_head_ was significantly reduced on SARS-CoV-2_live_-P17-IgG virions compared to controls, indicating P17-IgG-mediated S clustering. Boxplots showing the median (center line), interquartile range (box), and whiskers indicating the minimum and maximum values. Data points are excluded if the D_head_ is over 40 nm or less than 5 nm. (**G**) Proportion of S-trimers forming mode II(1) S-P17 Gemini complexes. On SARS-CoV-2_live_-P17-IgG, approximately 50% of prefusion S-trimers participated in mode II(1) complexes, whereas only ∼10% of S-trimers on SARS-CoV-2_fixed_-P17-IgG formed such complexes. Boxplots showing the median (center line), interquartile range (box), and whiskers indicating the minimum and maximum values. The boxplots in (F) and (G) highlight the time-dependent nature of Gemini complex formation on SARS-CoV-2_fixed_-P17-IgG.

To explore additional modes of bivalent binding, we first projected the S-P17 Solo complexes onto their refined coordinates on the viral surface. We then applied the same criteria used for identifying the S-P17 Gemini complex to determine whether two neighboring Fabs belonged to a single P17-IgG molecule. This analysis discovered ten additional S-P17 Gemini complexes modes , with their respective prevalence shown in Fig. 3C. These modes were categorized into two groups based on the number of P17-IgGs involved in bivalent interactions. In the first category, a single P17-IgG molecule coupled two S-trimers by engaging in any pairwise combinations among RBD1, RBD2 or RBD3. This category accounted for six S-P17 Gemini binding modes with a total prevalence of 23.4%. In the second category, two P17-IgG molecules cooperatively bridged two S-trimers, leading to the formation of five distinct S-P17 Gemini complexes, collectively representing 76.6% of the total structures. Among these, Mode II(1) was the most structurally dominant and was therefore characterized in detail (Fig. 2B). In the remaining mode II(2-5) complexes, the two S-trimers were coupled by P17-IgG molecules bridging various RBD pairs, including RBD3-RBD2, RBD2-RBD2, RBD3-RBD1, RBD2-RBD1, and RBD3-RBD3. Notably, in mode II(4) and mode II(5), one of the S-trimers exhibited a flipped orientation in which its up-RBD was positioned beneath the bridging IgGs, leading to greater separation of the S-P17 stems.

**Fig. 3.**
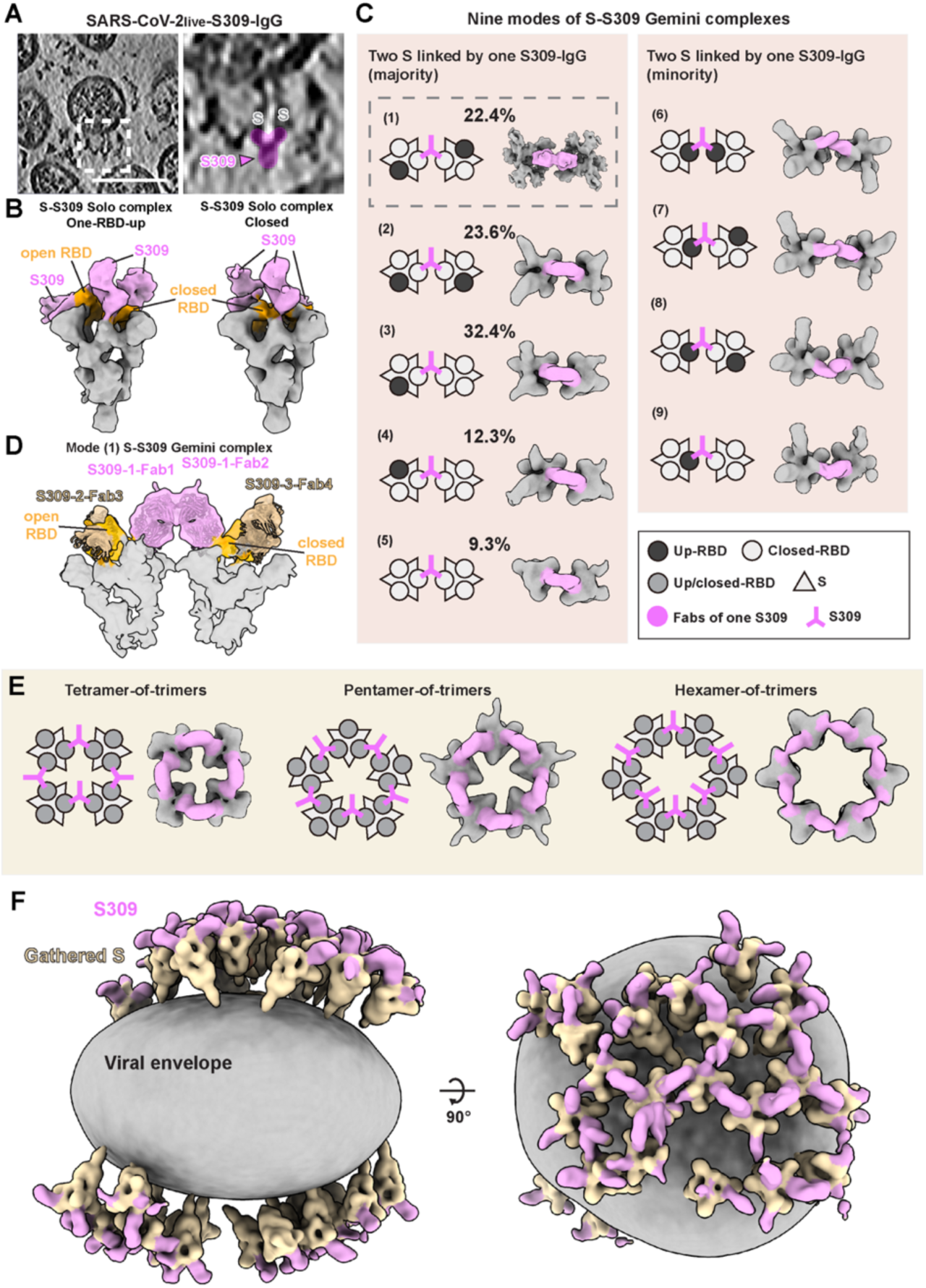
Bivalent binding of S309-IgG forms S-S309 Gemini complexes and gathers S-trimers into rings. (**A**) An exemplary tomogram slice (5 nm thickness) of SARS-CoV-2_live_-S309-IgG (30 minutes incubation) shows an S-S309 Gemini complex (white box), where two S-trimers are bridged by an IgG (pink). Scale bar: 100 nm. (**B**) Two conformations of S-S309 Solo complexes: one-RBD-up and closed conformations were resolved from SARS-CoV-2_live_-S309-IgG. RBD are colored orange and S309 are colored pink. (**C**) Schematics and their exemplary structures of the nine modes of S-S309 Gemini complexes, where one IgG bridge two S-trimers S. Modes (1-5) S-S309 Gemini complexes are coupled on their closed-RBDs and represent the major binding modes. Their structures and proportions were determined by STA. Modes (6-9) involve at least one up-RBD on their coupling sites and represent the minor binding modes. These complexes were identified by projecting the S-S309 Solo structures onto their refined coordinates on SARS-CoV-2_live_-S309-IgG. (**D**) Structure of mode (1) S-S309 Gemini complex. A single IgG molecule couples the RBDs of adjacent S-trimers, with RBD densities in orange, the bridging S309-IgG densities in pink and the other S309-IgG in tan. The structure was validated by fitting Fab densities with S309 Fab models from PDB: 6WS6. (**E**) Schematics and structures of the extended S-S309 multimers found on SARS-CoV-2_live_-S309-IgG, including a tetramer-, a pentamer-, and a hexamer-of S-trimers. The majority of multimers are circularly formed by S309-IgG bivalent binding. (**F**) A composite structure of a representative SARS-CoV-2_live_-S309-IgG virion was reconstructed by mapping all S-S309 complexes onto the viral envelope. S-trimers gathered into clusters are colored wheat. S309 are colored pink. The viral envelope (grey) was reconstructed by segmenting manually labeled densities.

Building on these modes, we found that P17-IgG actively gathered S-trimers and organized them into higher-order multimers (Fig. 2D). A frequently observed arrangement was a linear assembly of S-P17 complexes, where successive trimers are linked through bivalent IgG interactions. Although rare, trimer-of-trimers configurations were also detected. To visualize the complexity and collective impact of P17-mediated S-oligomerization on intact virions, we reconstructed SARS-CoV-2_live_-P17-IgG virions by mapping the S-P17 Solo and Gemini structures onto their refined coordinates. Compared to SARS-CoV-2_fixed_-apo^32^, the majority of S-trimers on SARS-CoV-2_live_-P17-IgG were gathered into patches, with interspersed exposed regions of the viral envelope (Fig. 2E). A similar gathering effect was also observed on SARS-CoV-2_fixed_-P17-IgG virions (fig. S3).

Assembling the mode II(1) Gemini complex requires precise spatial alignment of neighboring S-trimers at specific distances and angles. The observation that a substantial number of S-trimers successfully form such complexes on SARS-CoV-2_live_-P17-IgG is therefore unexpected. One possible explanation is that P17-IgG leverages its intrinsic flexibility and the mobility of S-trimers on the viral surface to facilitate S-trimer recruitment in assembling the mode II(1) complexes. To validate this, we first analyzed the inter-trimer distance between the head regions of the two closest S-trimers (D_head_) on virions. We observed a median D_head_ value of 14.2 nm for SARS-CoV-2_live_-P17-IgG across all incubation durations. In contrast, the same analysis conducted on SARS-CoV-2_fixed_-P17-Fab and SARS-CoV-2_fixed_-P17-IgG (15 minutes incubation) yielded median D_head_ values of 17.4 nm and 18.0 nm, respectively, which were comparable to the median value of 18.7 nm calculated from our prior SARS-CoV-2_fixed_-apo dataset^32^. After extended incubation, the D_head_ of SARS-CoV-2_fixed_-P17-IgG exhibited only a modest reduction to approximately 16.8 nm (Fig. 2F). These findings suggest that P17-IgGs facilitate rapid S-trimer clustering within minutes on live virions, whereas this clustering efficiency is markedly reduced on PFA-fixed virions, where S mobility and conformation are restricted. This phenomenon is further reflected in the proportion of mode II(1) Gemini complexes observed on virions. On SARS-CoV-2_live_-P17-IgG across all incubation durations, approximately 50% of prefusion S were incorporated into mode II(1) complexes, whereas only ∼10% of prefusion S were coupled into mode II(1) complexes on SARS-CoV-2_fixed_-P17-IgG (30 minutes incubation) (Fig. 2G). Collectively, these findings demonstrate that P17-IgG mediates diverse bivalent binding interactions that assemble S-trimers into clusters within minutes on live virions. The interplay between S-trimer mobility on the viral envelope and IgG flexibility appears to be crucial for the formation of these assemblies, which may enhance neutralization by increasing local antibody density and disrupting viral architecture.

### Bivalent Binding of S309-IgGs Forms Five Modes of S-S309 Gemini Complexes and Gathers S-trimers into Rings

To determine whether inter-spike bivalent binding and spike aggregation are unique to P17-IgG, we conducted analogous structural experiments using another anti-RBD monoclonal antibody, S309 (table S2). Although IC_50_ values of S309-IgG and S309-Fab were comparable in neutralizing SARS-CoV-2 pseudoviruses, only S309-IgG achieved 100% neutralization, indicating that IgG-specific bivalency plays a crucial role in viral inhibition^31^. WT SARS-CoV-2 were incubated with S309-IgG (2 μg/mL) at room temperature for 30 minutes, followed by PFA inactivation (SARS-CoV-2_live_-S309-IgG). Cryo-ET revealed distinct S309 densities localized at the head regions of the S-trimers (Fig. 3A). Preliminary STA and classification of 11,747 prefusion S from 492 SARS-CoV-2_live_-S309-IgG virions identified two major conformations of S-S309 complexes: a closed conformation (34.8%) and a one-RBD-up conformation (65.2%) (figs. S5A and S6A). At lower thresholds, emerging densities extending from each RBD were observed. Notably, a single closed-RBD exhibited the strongest density across both S-S309 complexes, forming a secondary S-S309 complex. This suggests that S309- IgG may also engage in inter-spike bivalent binding on-virion. Indeed, further classification revealed two S-S309 Solo complexes (Fig. 3B and fig. S4) and five distinct S-S309 Gemini complexes. Additionally, by mapping the S-S309 Solo complexes onto their refined coordinates, we identified four additional S-S309 Gemini complexes (Fig. 3, C and D). The two S-S309 Solo complexes—one in a one-RBD-up and the other in a closed conformation— demonstrated S309-IgG binding to all RBDs (Fig. 3B and fig. S4B). These structures closely aligned with their respective recombinant S-S309-Fab structures (PDB: 6WPT, 6WPS)^31^ (fig. S4C). On the one-RBD-up S-S309 Solo complex, Fab densities on the closed-RBDs were relatively stronger than on the up-RBD, whereas only one Fab density was distinctly present in the closed S-S309 Solo complex (Fig. 3B). We further quantified the presence of postfusion S, revealing that 6% S-trimers on SARS-CoV-2_live_-S309-IgG were in postfusion conformation.

Unlike P17-IgG, which frequently forms double-bridged Gemini complexes, S309-IgG exclusively assembles single-bridged Gemini complexes. Among the nine modes of S-S309 Gemini complexes observed on-virion, the majority were associated with the closed-RBDs of adjacent S-trimers; while up-RBDs, although less frequent, also contributed to S-coupling (Fig. 3C). The distribution of S-S309 Gemini complex modes is relatively balanced, in contrast to the predominant mode II(1) in S-P17 Gemini complexes. Notably, 91% of S-S309 Gemini complexes incorporate at least one one-RBD-up S. Together with the relatively weaker Fab densities observed on closed S, our findings suggest that S309-IgG preferentially binds to the one-RBD-up S. Further mapping of all S-S309 Solo and Gemini complexes onto their refined coordinates revealed that S309-IgG induces distinct spatial arrangements of S-trimers compared to P17-IgG. Specifically, S309-IgG preferentially organizes S-trimers into ring-like assemblies, including tetramer-, pentamer-, and hexamer-of-trimers (Fig. 3E and fig. S5). Within these ring structures, each S-trimer is linked to its two neighboring S-trimers via S309-IgGs. This coupling mechanism may facilitate the formation of larger multimers, such as heptamer- or octamer-of-trimers. Furthermore, bivalent binding by S309-IgG also leads to the formation of multi-ring lattice structures on virions (Fig. 3F). This spatial organization markedly contrasts with the linear assemblies of S-trimers induced by P17-IgG.

Collectively, these findings demonstrate that although both antibodies exploit inter-spike bivalent binding to cluster S-trimers, S309-IgG preferentially organizes S-trimers into circular arrangements due to differences in epitope accessibility and binding orientation. These differences likely arise from the distinct structural properties of the S309 and P17 epitopes, as well as their influence on S-trimer conformations upon binding. S309-IgG’s ability to assemble S-trimers into rings and lattices highlights the versatility of bivalent binding in shaping viral surface architecture, providing further insights into the structural basis of IgG-mediated neutralization.

### IgGs Can Aggregate Virions Through Inter-Virion Bivalent Binding

Cryo-ET analysis of SARS-CoV-2 virions incubated with P17-IgG or S309-IgG revealed occasional virion aggregation, suggesting that IgGs may facilitate inter-virion coupling through bivalent binding. Examination of SARS-CoV-2_live_-P17-IgG tomogram identified densities corresponding to S-trimers from neighboring virions connected by a single P17-IgG molecule (Fig. 4A). By mapping the S-P17 Solo and Gemini complexes onto their refined coordinates, we confirmed that the observed inter-virion coupling resulted from bivalent interactions of a single P17-IgG engaging RBDs on adjacent virions (Fig. 4B). To assess the functional impact of IgG-induced virion aggregation, we performed dynamic light scattering (DLS) experiments using SARS-CoV-2_fixed_-apo. Over a two-hour incubation period, the mean hydrodynamic radius of the sample increased progressively, consistent with virion clustering. In contrast, no significant increase in particle size was observed in control samples lacking P17-IgG, confirming that the aggregation was IgG-dependent (fig. S6). A similar phenomenon was observed in SARS-CoV-2_live_-S309-IgG tomograms, where coupling occurred through S309-IgG-mediated bivalent interactions. Composite reconstruction of these virions revealed a pentamer-of-trimers inter-virion complex containing four S-trimers from one virion and one S-trimer from a neighboring virion. This assembly was stabilized by multiple S309-IgG molecules bridging the virions (Fig. 4C). To quantify the relative contributions of inter- and intra-virion bivalent binding, we analyzed the proportion of inter- and intra-virion bivalent binding events between adjacent S-trimers. Statistical analysis revealed that inter-virion binding accounted for only ∼5% of all bivalent interactions, while intra-virion binding remained the predominant mode of IgG engagement (Fig. 4D). Despite its lower prevalence, inter-virion binding highlights the ability of IgGs to promote virion aggregation under specific conditions, potentially enhancing immune recognition and clearance.

**Fig. 4.**
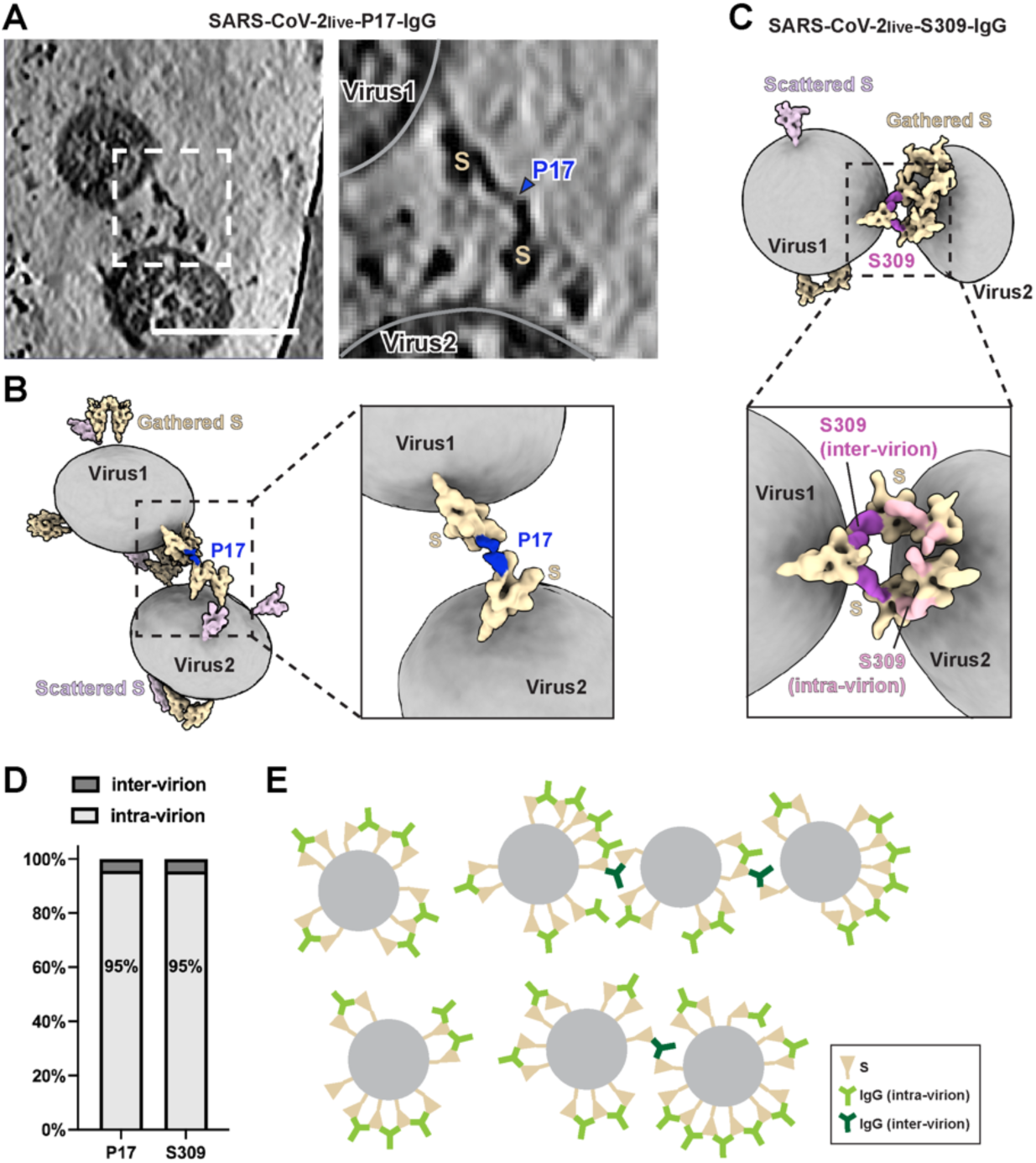
Bivalent binding by IgGs can induce inter-virion aggregation. (**A**) An exemplary tomogram slice (5 nm thickness) highlights an S-P17 complex (white box), where a single P17-IgG bridges two opposing S-trimers from two neighboring SARS-CoV-2_live_-P17-IgG virions (15 min incubation). Scale bar: 100 nm. (**B**) A composite reconstruction of the virions in **a** illustrates how a single P17-IgG molecule (colored dark blue) connects S-trimers (wheat) on neighboring virions (gray viral envelopes). Scattered S-trimers are shown in thistle. A close-up of the inter-virion S-P17 complex (black box) highlights the bridging interaction. (**C**) A composite reconstruction of a SARS-CoV-2_live_-S309-IgG shows an S309-IgG pentamer-of-trimers, where four S-trimers from one virion and one S-trimer from a neighboring virion are coupled by two S309-IgG molecules (purple). Viral envelopes (gray), gathered S-trimers (wheat), and scattered S-trimers (thistle) are visualized, with a close-up of the inter-virion complex (black box). (**D**) Quantitative analysis revealed that inter-virion bivalent binding accounts for ∼5-6% of total bivalent binding events in both SARS-CoV-2_live_-P17-IgG and SARS-CoV-2_live_-S309-IgG samples. Intra-virion binding remains the predominant mode of IgG-mediated interactions. (**E**) A schematic illustrates the collective effects of IgG molecules in organizing S-trimers within a single virion (intra-virion bivalent binding, green) and couple S-trimers on neighboring virions (inter-virion bivalent binding, dark blue).

Based on these findings, we proposed a model outlining the collective effects of IgG bivalent binding in viral organization (Fig. 4E). Intra-virion binding organizes S-trimers into higher-order assemblies on individual virions, while inter-virion binding enables IgGs to link neighboring virions into clusters. This dual capability underscores the versatility of IgG in modulating viral surface architecture and promoting aggregation.

## Discussion

In this study, we employed cryo-ET to characterize the structural mechanisms by which two potent anti-RBD nAbs, P17 and S309, engage with S-trimers on SARS-CoV-2. Our findings demonstrate that intact IgGs engage in distinct oligomerization patterns compared to their Fab counterparts, forming inter-spike and inter-virion interactions that significantly alter the structural landscape of the viral surface. The structures of on-virion S-Fab and S-IgG Solo complexes largely recapitulate their corresponding recombinant S-Fab structures. However, the Gemini and ring structures of S-IgG, as well as the lack of observed two-RBD-up S-P17-Fab structures (PDB: 7CWL) on-virion, highlight key differences between in situ and recombinant structural studies. Several factors may contribute to these discrepancies. First, intact IgGs were used in our study, whereas prior recombinant studies utilized Fabs. Second, prefusion S in their native virion context and those used for recombinant protein studies often exhibit different conformational distributions. For example, on-virion S-trimers predominantly adopt in a closed or one-RBD-up state at an approximate ratio of 1:1^32^. In contrast, prefusion-stabilized ectodomain S that exclusively adopts a one-RBD-up conformation^33^ were used for recombinant studies^30^. Furthermore, our findings emphasize the role of spatial organization in IgG-mediated viral inhibition. Unlike recombinant proteins in solution, which rely on Brownian motion and concentration gradients for inter-spike binding, on-virion S-trimers are constrained by the membrane. Their spatial distribution, swinging^32^ and diffusion^34,35^ features on the viral envelope, together with the IgG flexibility, facilitate rapid and stable multimerization of S-trimers. This explains why the tentative inter-spike bivalent S-IgG structures determined from recombinant proteins are often in a head-to-head configuration^36,37^, different from the Gemini and ring structures we reported.

Next, we aim to rationalize the differences between P17 and S309 in their capacity to form S-IgG multimers. While both antibodies bind to RBDs in either the up- or closed-conformations, they exhibit distinct modes of S-IgG multimerization. These differences stem from variations in their epitopes and their respective impacts on S-trimer conformation upon binding. On the viral surface, P17 stabilizes all prefusion S in a one-RBD-up conformation, resulting in the movement of P17-1 to P17-2 (Fig. 5A). In category I S-P17 Gemini complexes, a single P17-IgG bridges two S-trimers, minimizing steric clashes. Conversely, in category II, two P17-IgGs couple S-trimers, necessitating spatial adjustments to avoid steric hindrance (Fig. 5B). This is particularly evident in modes II(2-5), where the two S-trimers adopt unfavorable spatial arrangements, requiring increased spacing between their stalk regions to prevent clashes at the N-terminal domains (NTD). Notably, Mode II(1) is an exception, representing the predominant structural population of prefusion S on P17-treated virions and the only structurally resolved Gemini complex. This implies that Mode II(1) achieves an optimal stability likely due to cooperative structural interactions at both the head and stalk regions of the S-trimers. At the head region, the two S1 subunits are symmetrically coupled via their RBD1-RBD2 regions by a pair of P17-bridges, reinforcing the Gemini structure. At the stalk region, the two stalks maintain close alignment and remain nearly parallel. In comparison, S-S309 Gemini complexes are exclusively coupled by a single S309-IgG. The even distribution of S309-IgGs across the head regions of S-trimers, regardless of the one-RBD-up or closed conformational states (Fig. 5C), precludes the formation of double-bridged Gemini complexes due to steric clashes at the NTDs (Fig. 5D). Additionally, the distinct epitopes of P17 and S309 influence how their respective S-IgG Gemini complexes extend into higher-order multimers. P17 preferentially forms linear S-assemblies by linking successive S-P17 Gemini complexes (Fig. 5E), whereas S309 more frequently drives circular S-assemblies, which can further expand into lattice-like structures (Fig. 5F).

**Fig. 5.**
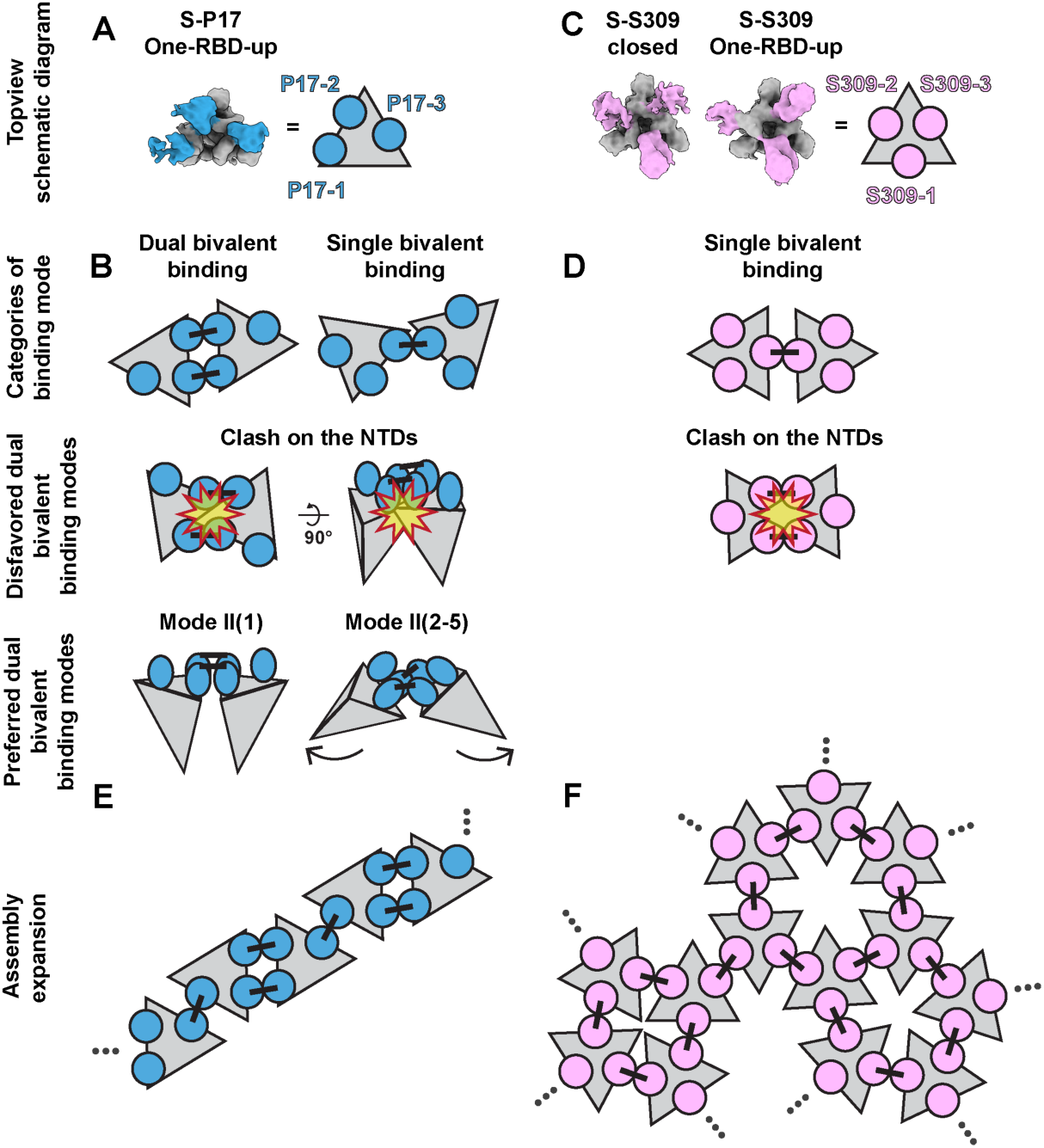
Comparison of P17 and S309 bivalent binding and assembly mechanisms. Schematic representation illustrates how differences in epitope binding orientations and IgG’s impact on S-trimer’s conformations lead to distinct patterns of S-IgG assemblies. (**A**) P17-IgG (blue circles) stabilizes S-trimers in a one-RBD-up conformation, forming two neighboring P17-IgGs on one side of S-trimer’s head region (grey triangle) and a P17-IgG on the other side. (**B**) This configuration facilitates the formation of both single- and double-bridged Gemini complexes. However, the steric clashes on NTD only allow the configuration of mode II(1) Gemini, which has parallel S-trimer stems, or modes II(2-5), which have separated stems. (**C**) S-S309 complexes exhibit both closed and one-RBD-up conformations. S309-IgGs (pink circles) are uniformly distributed across the head regions of S-trimers in both conformations. (**D**) This configuration allows the formation of single-bridged Gemini complexes but inhibits the double-bridged ones. (**E**) S-P17 complexes are often extended into linear assemblies by P17-IgG. (**F**) S-S309 complexes are often extended into circular and lattice assemblies by S309-IgG.

IgG-mediated immunity against enveloped viruses operates primarily through two mechanisms: neutralization and Fc-mediated effector functions^11^. Our study provides structural insights into IgG avidity and offers novel evidence supporting these immunological pathways. For neutralization, we demonstrate that P17-IgG exhibits enhanced affinity through bivalent binding compared to P17-Fab, enabling more effective epitope engagement on virions. Additionally, the avidity interactions between IgG and the S-trimer can induce S1 shedding. Through versatile modes of bivalent binding, both P17 and S309 effectively gather prefusion S-trimers on the viral surface. This process likely introduces tension within the S1 subunits, leading to their shedding, as evidenced by our cryo-ET and Western blot analyses. This mechanism may explain why certain IgGs exhibit greater potency in inducing S1 shedding compared to their Fab counterparts^9^. Notably, compared to P17-Fab, P17-IgG did not demonstrate significantly higher neutralization potency against live SARS-CoV-2. This outcome may be attributed to the potent neutralization efficiency of P17-Fab, thereby limiting the additional enhancement conferred by IgG bivalency. In terms of Fc-mediated effector functions, the multimeric forms of S-P17 and S-S309 generate diverse patterns of , which display enhanced Fc-avidity for multivalent interactions with FcRs and may promote Fc-mediated immune responses^10^. For instance, the pentameric and hexameric S-S309 multimers could serve as structural platforms for complement pathway activation^38^. Moreover, it is plausible that P17 and S309 assemble similar S-IgG complexes on the membranes of SARS-CoV-2-infected cells, thereby triggering cell-associated immune responses such as antibody-dependent cellular cytotoxicity (ADCC), antibody-dependent cellular phagocytosis (ADCP), and antibody-dependent complement activation. Notably, these effects have already been confirmed for S309^39^.

Another compelling observation is the ability of IgG to mediate inter-virion coupling through bivalent binding. Although inter-virion bivalent binding constitutes only 5-6% of total bivalent interactions, its occurrence suggests an additional mechanism by which IgGs influence viral dynamics. By linking adjacent virions into clusters, IgG-mediated aggregation may enhance immune recognition and activate effector functions, thereby contributing to overall antiviral efficacy. However, the physiological relevance of this phenomenon remains uncertain. In vivo conditions may impose constraints on IgG-induced aggregation due to lower viral loads and antibody concentrations. In our experiments, when live SARS-CoV-2 virions were incubated with IgGs to generate the SARS-CoV-2_live_-P17-IgG and SARS-CoV-2_live_-S309-IgG samples, the final concentrations of virions and IgGs were approximately 10⁷ virions/mL and 2 μg/mL, respectively. By comparison, the peak viral load of WT SARS-CoV-2 detected in nasal samples is approximately 10^5^ virions/mL^40^, and a comparable RBD-specific IgG concentration of approximately 10 μg/mL has been reported in patients’ serum^41^. While some studies suggest that IgG can aggregate SARS-CoV-2 via crosslinking^36,37^, our data indicate that such aggregation events are relatively uncommon in vivo, except in regions with exceptionally high localized viral concentrations.

The significant variation in avidity among nAbs targeting enveloped viruses^42^ highlights the necessity of understanding how IgGs achieve bivalency at the molecular level, taking into account antibody structure, epitope accessibility, and antigenic organization. Several key factors contribute to this process. First, the Fab association rate constant influences bivalent binding efficiency. Upon initial Fab binding to an epitope, a restricted reaction volume enhances the probability of the free Fab arm binding to an adjacent epitope. IgGs with a higher association rate constant in their half-bound state are more likely to engage in efficient bivalent interactions^43^. Second, bivalent binding is largely influenced by epitope accessibility and conformational independence. Epitopes located on membrane-distal regions of glycoproteins are more conducive to bivalent binding than those near membrane-proximal stalks, where steric hindrance is more pronounced^27,44^. Moreover, epitopes that remain structurally accessible regardless of glycoprotein conformation, such as the class-2 and class-3 RBD-nAbs^29^ or 101F epitope of the RSV F protein^45^, are more likely to support bivalent interactions. Third, the orientation of Fab binding influences whether IgGs engage in intra- or inter-spike bivalency. For example, P17 and S309 Fabs bind all three epitopes of an S-trimer with their C-termini pointing outward, facilitating inter-spike bivalent interactions. In contrast, IgGs with downward-facing Fab orientations targeting S-trimers are less likely to participate in inter-spike bivalent binding due to steric interference from their Fc domains^7^. Finally, inter-spike bivalency is modulated by the antigen landscape on the viral surface. Factors such as glycoprotein molecular mass, oligomerization state, density, and mobility play a crucial role in shaping IgG binding modes. Viruses with densely packed glycoproteins, such as influenza and Lassa viruses^14^, or those with large and highly mobile spikes, such as SARS-CoV-2, are more susceptible to inter-spike bivalent binding compared to viruses with sparse or relatively static glycoproteins^46^.

Our findings also have implications for natural antibody screening, therapeutic antibody design and vaccine development. The comparison of bivalent binding efficiency on SARS-CoV-2_live_-P17-IgG and SARS-CoV-2_fixed_-P17-IgG samples has led us evaluate the potential effectiveness of chemically crosslinked SARS-CoV-2 as vaccines. We demonstrated that chemical crosslinking deprives S-trimers of their mobility and conformational dynamics on fixed SARS-CoV-2, resulting in significantly reduced avid interactions with antibodies compared to those on live SARS-CoV-2. This limitation likely impacts B cell activation and the subsequent production of nAbs^47^. In addition, the ability of P17 and S309 to induce distinct S-trimer assemblies suggests that optimizing epitope presentation, oligomerization and accessibility on vaccine platforms could enhance immunogenicity. Overall, our structural analysis of IgG avidity highlights the importance of antigen mobility and multimerization in eliciting effective immune responses. These findings inspire further investigations into the multivalent interactions between viruses and various antibody types, including IgG, IgA, IgM, and engineered multimeric antibodies.

## Materials and Methods

### Cells, virus and antibody

Vero cells (ATCC CCL-81) were cultured in Modified Eagle Medium (MEM; Corning Inc., Corning, NY) supplemented with 10% fetal bovine serum (FBS; Gibco, Carlsbad, CA), Penicillin (100 U/mL) and streptomycin (100 mg/mL) (Gibco, Carlsbad, CA). SARS-CoV-2 virions were propagated using Vero cells as described^32^. On the fourth day post-infection, the cell supernatant was collected, cleared of cell debris by centrifugation at 4,000 g for 30 min. Titration of virus in the cell supernatant was measured by TCID_50_ assay. The virus solution was diluted in a 10-fold gradient series in MEM culture medium containing 2% FBS. 100 μL of diluted virus solution per well was added to a 96-well plate (Greiner Bio-One, Germany). MEM medium without virus solution was used as normal cell control. Then, 10^4^ cell/100 μL Vero cells were added in each well, and incubated in 5% CO_2_ at 35°C. The cytopathic effect (CPE) was observed and recorded on day 5 of culture. According to the CPE results, the TCID_50_ of the virus was calculated by Spearman-Karber method. There were 8 wells in each group, and the experiment was repeated for 3 times. The measured infectious viral titer is 10^7.2^ TCID_50_/mL.

For the SARS-CoV-2_fixed_-apo sample, the virus-containing supernatant was cleared of cell debris and inactivated with paraformaldehyde (PFA; final concentration 3%) for 48 h in 4℃. For the SARS-CoV-2_live_-P17-IgG sample, the debris-cleared supernatant was incubated with monoclonal antibody P17-IgG (2 μg/mL) at room temperature for 15 min, 30 min, 1 h, 2 h or 4 h, followed by inactivation with PFA. For the SARS-CoV-2_live_-P17-Fab sample, the debris-cleared supernatant was incubated with P17-Fab at a concentration of 1.33 μg/mL, room temperature for 15 min, followed by inactivation with PFA. For the SARS-CoV-2_fixed_-P17-IgG sample, the SARS-CoV-2_fixed_-apo was purified through a 30% sucrose cushion by ultracentrifugation at 100,000 g, 4℃ for 3 h, then resuspended in PBS buffer and incubated with P17-IgG (43.5 µg/mL) at room temperature for 15 min, 30 min, 1 h or 2 h. For the SARS-CoV-2_live_-S309-IgG sample, the debris-cleared supernatant was incubated with monoclonal antibody S309-IgG (2 μg/mL) at room temperature for 30 min, followed by inactivation with PFA.

All the experiments involving infectious SARS-CoV-2 were carried out in an approved biosafety level (BSL)-3 laboratory of Zhejiang University. Purification, concentration and EM sample preparation of fixed virus were conducted in a BSL-2 lab at Tsinghua University.

### Antibody expression and identification

Expression and purification of P17-IgG antibody was performed as described^30^. Briefly, plasmids harboring the heavy-chain and light-chain of P17 were co-transfected into ExpiCHO cells. The cell supernatant was collected and purified by MabSelect SuRe LX (Cytiva, Marlborough, MA) after 7 to 15 days of transfection.

P17-Fab antibody was expressed using the ExpiCHO system (Thermo Fisher Scientific). Eukaryotic expression plasmid pcDNA3.3-TOPO (Invitrogen) containing separate heavy-chain VH and CH1, light-chain VL and CL1 were co-transfected into ExpiCHO cells. After 7 to 15 days of transfection, the cell supernatant was collected and purified by Ni-NTA (Thermo Fisher Scientific). Finally, the purified antibody was exchanged into PBS buffer and concentrated through the centrifugal filter unit (Millipore).

The neutralization efficiency of P17-IgG and P17-Fab against SARS-CoV-2 was measured by virus microneutralization assay. In brief, antibodies were diluted in a two-fold series, combined with 100 TCID_50_ of SARS-CoV-2 viruses, and then the mixture was incubated for 2 h at 35℃. After that, Vero cells were added and incubated at 35℃ with 5% CO_2_. The neutralization efficiency was assessed by observing CPE on day 5 of the culture, thereby confirming the assay outcome.

Plasmids encoding the heavy and light chains of S309-IgG were transiently co-transfected into HEK293F cells using PEI MAX (Polysciences) in a molecular ratio of 1:2. After 4 days of transfection, the cell supernatant containing S309-IgG were collected and centrifugated at 4,000 r.p.m. for 20 min to remove cell debris. S309-IgG in the supernatant were purified by a Hitrap Protein A HP column (Cytiva) and gel filtration chromatography using Superdex 200 Increase 10/300 GL (Cytiva).

### Cryo-ET sample preparation

For the SARS-CoV-2_live_-P17-IgG, SARS-CoV-2_live_-P17-Fab and SARS-CoV-2_live_-S309-IgG samples, virions were purified through a 30% sucrose cushion by ultracentrifugation at 100,000 g, 4℃ for 3 h, then resuspended in PBS buffer. An 8 μL aliquot of the concentrated virus sample was applied onto a glow-discharged copper grid coated with holey carbon film (R2/2, 300 mesh, Quantifoil, Jena, Germany). Grid was skimmed over PBS to remove residual sucrose. After adding 3 μL of 10-nm gold tracer (Aurion, the Netherlands), the grid was blotted for 4S-trimerand plunge-frozen into liquid ethane using a cryo-plunger 3 (Gatan Inc., CA). For the SARS-CoV-2_fixed_-P17-IgG sample, the virus was applied directly onto the grids, which were then plunge-frozen in the same manner as described above.

### Cryo-ET data acquisition

Grids were imaged on a Titan Krios microscope (Thermo Fisher Scientific, Hillsboro, OR) operated at 300 kV equipped with a K3 direct electron detector (Gatan Inc., CA) and a BioQuantum energy filter (slit width 20 eV). Virions were recorded in super-resolution mode at a nominal magnification of 64,000×, resulting in a calibrated pixel size of 0.68 Å. Tilt series were recorded with eight or ten frames at each angle using a total dose of 131.2 e^-^/Å^2^ and defoci ranging from -2 to -4 μm in SerialEM^48^. For SARS-CoV-2_live_-P17-IgG sample, 169 sets of tilt series using dose-symmetric scheme from -60° to 60° and 128 using bidirectional scheme from 15° to -60°, then 18° to 60° at 3° steps were collected. For SARS-CoV-2_live_-P17-Fab sample, 54 sets of tilt series using dose-symmetric scheme were collected. For SARS-CoV-2_fixed_-P17-IgG sample, 65 sets of tilt series using bidirectional scheme were collected. For SARS-CoV-2_live_-S309-IgG sample, 38 sets of tilt series using dose-symmetric scheme were collected.

### Cryo-ET data processing

Tilt series data were preprocessed using a high-throughput suite^32^ developed in our lab. Motion between frames were corrected using MotionCor^49^. Cumulative exposure was filtered as described^50^. Defocus values were estimated using GCTF^51^. Tilt series were aligned by tracking gold fiducials using IMOD^52^. 276 tomograms of the SARS-CoV-2_live_-P17-IgG sample, 60 tomograms of the SARS-CoV-2_fixed_-P17-IgG sample, 54 tomograms of the SARS-CoV-2_live_-P17-Fab sample and 33 tomograms of the SARS-CoV-2_live_-S309-IgG sample with good alignment were reconstructed by weighted-back projection after correcting the contrast transfer function (CTF) using NovaCTF^53^, resulting in a final pixel size of 1.36 Å/pixel. For particle picking, tomograms were either 4 × binned and low-pass filtered to 80 Å, or 8 × binned and denoised using cryoCARE^54^. S-trimers were manually picked and their initial orientations were given from vectors normal to the envelope. For improved visualization and presentation, tomograms were 8 × binned and missing-wedge corrected using IsoNet^55^. Subtomogram averaging (STA) was carried out using Dynamo^56^.

### Structure fitting

To interpret the S-P17 Solo structure, the recombinant model of the one-RBD-up S-P17-Fab (PDB: 7CWM) was fitted into our structure using ChimeraX. Since the constant domain of P17 Fab was not included in the recombinant S-P17 model, models of an intact class-2 nAb Fab (PDB: 7K8O) were subsequently fitted to the Fabs densities of the S-P17 Solo structure. To interpret the mode II(1) S-P17 Gemini structure, two recombinant models of the one-RBD-up S-P17-Fab (PDB: 7CWM) and six models of Fab (PDB: 7K8O) were fitted into the structure using ChimeraX. Similarly, the recombinant models of the S-S309-Fab in the one-RBD up and closed conformation (PDB: 6WPT, 6WPS), and the models of the complete S309-Fab (PDB: 6WS6) were fitted into the S-S309 Solo and Gemini complex structures using ChimeraX.

To determine whether two adjacent Fabs belong to a P17-IgG or S309-IgG molecule in S-P17 and S-S309 complexes, the distance between the heavy chain residues 222 of the fitted 7K8O and 6WS6 models was measured. If the distance was less than 65 Å and the C termini of adjacent Fabs were in an inward-pointing orientation, the two Fabs were regarded as part of a P17-IgG or S309-IgG molecule^57^.

### Western blotting

A plasmid encoding SARS-CoV-2 WT strain (Wuhan-Hu-1) S protein was transfected into HEK293T cells using the Lipo8000 transfection reagent (Beyotime Biotec. Inc., Shanghai, China). At 24 hours post-transfection, HEK293T cells were removed from the plate with 0.2 mg/mL EDTA and washed twice with 100 μL ice-cold PBS (Gibco, Carlsbad, CA) containing 2% FBS (Gibco, Carlsbad, CA). Approximately 10^5^ cells per sample were incubated in the presence or absence of 2 μg/mL P17 in 100 μL PBS containing 2% FBS for 0, 15, 30, 60, 120 or 240 min at 37°C. After the allocated incubation periods, samples were immediately placed on ice, and cell debris was removed by centrifugation. SDS-PAGE sample Loading Buffer (Beyotime Biotec. Inc., Shanghai, China) was added, and the mixture was boiled at 100°C for 20 min. All samples were run on a 4-20% gradient Bis-Tris gel (GenScript Biotech Corporation, Jiangsu, China) and transferred to a polyvinylidene fluoride (PVDF) membrane (Merck Millipore Ltd., Co. Cork, Ireland) at a constant voltage of 20 V for 35 min. Total protein normalization was performed on the transferred PVDF membrane using the No-Stain Protein Labeling Reagent (Invitrogen, Carlsbad, CA). The membrane was blocked with 6% (w/v) non-fat milk (Sangon Biotech Co., Shanghai, China) diluted in PBS containing 0.1% Tween-20 (PBS-T) for 60 minutes at room temperature. Anti-SARS-CoV-2 Spike S1 NTD monoclonal antibody (Cell Signaling Technology, MA; 1:1000 dilution), anti-SARS-CoV-2 Spike S2 polyclonal antibody (Sino Biological Inc., Beijing, China; 1:4000 dilution), Horse Radish Peroxidase (HRP)-conjugated goat anti-mouse IgG secondary antibody (Sino Biological Inc., Beijing, China; 1:5000 dilution) and HRP-conjugated goat anti-rabbit IgG-Fc secondary antibody (Sino Biological Inc., Beijing, China; 1:10000 dilution) diluted in PBS-T were used for Western blotting. HRP detection was performed using an SuperSignal West Pico PLUS Chemiluminescent Substrate (Thermo Fisher Scientific, Bremen, Germany) and imaged with a GelView 6000Plus Imager (BLT photon technology, Guangdong, China). Western blot bands were quantified using ImageJ. Statistical analysis was performed with GraphPad Prism version.

### Dynamic light scattering

The SARS-CoV-2_fixed_-apo sample was purified through a 30% sucrose cushion by ultracentrifugation at 100,000 g, 4℃ for 3 h, resuspended in PBS buffer and mixed with or without P17-IgG (final concentration 100 μg/mL) at 25℃. Dynamic light scattering measurements were taken immediately after incubation at 10-minute intervals and continued for 2 h on a DynaPro Nanostar instrument (Wyatt Technology, CA). 30 measurements were taken at each time point to obtain the radius distribution of solution.

### Quantification and statistical analysis

All data was analyzed and plotted using GraphPad Prism. For the pre- and postfusion S count per virion (Fig. 1e), 2,020 SARS-CoV-2_fixed_-apo virions^32^, 72 SARS-CoV-2_live_-P17-Fab virions, 78 SARS-CoV-2_live_-P17-IgG virions (15 minutes incubation), 79 SARS-CoV-2_fixed_-P17-IgG virions (15 minutes incubation), 75 SARS-CoV-2_live_-P17-IgG virions (30 minutes incubation), 73 SARS-CoV-2_fixed_-P17-IgG virions (30 minutes incubation), 74 SARS-CoV-2_live_-P17-IgG virions (1 hour incubation), 74 SARS-CoV-2_fixed_-P17-IgG virions (1 hour incubation), 69 SARS-CoV-2_live_-P17-IgG virions (2 hours incubation), 71 SARS-CoV-2_fixed_-P17-IgG virions (2 hours incubation), and 69 SARS-CoV-2_live_-P17-IgG virions (4 hours incubation) were used for analysis.

For the distance between the head regions of two nearest S-trimers on each P17-incubated SARS-CoV-2 virion (D_head_, Fig. 2f), the D_head_ was calculated according to the refined S coordinates from subtomogram averaging. The defined S head region is 76 Å above the center of the resulting subtomogram averaging map of S-P17 Solo complexes, near the top of S and the center of residues 503 in RBD-down conformation. 52,380 prefusion S from 2,020 SARS-CoV-2_fixed_-apo virions^32^, 12,068 prefusion S from 599 SARS-CoV-2_live_-P17-Fab virions, 20,486 prefusion S from 1,611 SARS-CoV-2_live_-P17-IgG virions (15 minutes incubation), 6,197 prefusion S from 599 SARS-CoV-2_live_-P17-IgG virions (30 minutes incubation), 5,458 prefusion S from 590 SARS-CoV-2_live_-P17-IgG virions (1 hour incubation), 2,732 prefusion S from 284 SARS-CoV-2_live_-P17-IgG virions (2 hours incubation), 1,058 prefusion S from 171 SARS-CoV-2_live_-P17-IgG virions (4 hours incubation), 1,826 prefusion S from 192 SARS-CoV-2_fixed_-P17-IgG virions (15 min incubation), 2,906 prefusion S from 207 SARS-CoV-2_fixed_-P17-IgG virions (30 min incubation), 2,084 prefusion S from 140 SARS-CoV-2_fixed_-P17-IgG virions (1 hour incubation), and 3,709 prefusion S from 346 SARS-CoV-2_fixed_-P17-IgG virions (2 hours incubation) were used for analysis. Distances greater than 40 nm or less than 5 nm were excluded from the analysis.

As to the proportion of S that composes S-P17 Gemini complexes in prefusion S per virion (Fig. 2g), 9,835 S-P17 Gemini particles and 18,596 S-P17 Solo complexes particles on 3,255 SARS-CoV-2_live_-P17-IgG virions were used for statistics. 1,611 virions (15 minutes incubation), 599 virions (30 minutes incubation), 590 virions (1 hour incubation), 284 virions (2 hours incubation) and 171 virions (4 hours incubation) were used, respectively. For the SARS-CoV-2_fixed_-P17-IgG sample, 818 S-P17 Gemini complexes, 4,884 S-P17 Solo complexes in one-RBD-up conformation and 4,628 S-P17 Solo complexes in closed conformation on 885 SARS-CoV-2_fixed_-P17-IgG virions were used for statistics. 192 virions (15 minutes incubation), 207 virions (30 minutes incubation), 140 virions (1 hour incubation) and 346 virions (2 hours incubation) were used respectively. Each S-P17 Gemini complex contains two S-trimers.

For the ratios of inter-virion and intra-virion bivalent binding (Fig. 4d), IgG bivalent binding interactions was assessed using the 65 Å distance criterion between Fabs. The coordinates of 14,006 S-P17 Solo complexes and 7,340 S-P17 Gemini complexes from 2,210 SARS-CoV-2_live_-P17-IgG virions (15 minutes and 30 minutes incubation) were analyzed for Fab distance measurements. A total of 10,917 bivalent binding interactions were observed, including 10,433 intra-virion bivalent binding interactions and 484 inter-virion bivalent binding interactions. For SARS-CoV-2_live_-S309-IgG, the coordinates of the one-RBD-up and closed S-S309 complex (prior to classification as Solo or Gemini structures) were analyzed for Fab distance measurements. Distances from 11,746 prefusion S on 492 SARS-CoV-2_live_-S309-IgG virions were measured, resulting in 6,131 bivalent binding interactions, including 5,850 intra-virion bivalent binding interactions and 281 inter-virion bivalent binding interactions.

For the ratios of S-P17 Gemini complex (Fig. 2c), IgG bivalent binding interactions were assessed using the 65 Å distance criterion between Fabs. The coordinates of 14,006 S-P17 Solo complex and 7,340 S-P17 Gemini complex from 2,210 SARS-CoV-2_live_-P17-IgG virions (15 minutes and 30 minutes incubation) were analyzed for Fab distance measurements. 2,382 category I bivalent binding interactions and 7,694 category II bivalent binding interactions were determined from the intra-virion bivalent binding interactions. In the category I, 154 mode I(1), 164 mode I(2), 325 mode I(3), 527 mode I(4), 652 mode I(5) and 560 mode I(6) bivalent binding interactions were determined. In the category II, 7,340 mode II(1), 49 mode II(2), 144 mode II(3), 44 mode II(4), 23 mode II(5) and 35 other forms of bivalent binding interactions were determined.

## Acknowledgements

We thank Dr. Jianlin Lei, Dr. Fan Yang and Dr. Xiaomin Li from the cryo-EM Facility, Technology Center for Protein Sciences, Tsinghua University, for their support on cryo-EM data collection. We thank the computational facility support on the cluster of Bio-Computing Platform (Tsinghua University Branch of China National Center for Protein Sciences Beijing).

## Declarations

### Author contributions

S.L. and H.Y. conceived and supervised the project. H.Y., M.Z., Q.H., X.Z., D.S., Z.W., X.L. and H.W. provided the virus, antibody, antibody-incubated virus samples and analyzed the antiviral activity of antibodies. Y.S. and Q.H. isolated the viruses, performed biochemical analysis and prepared the cryo-sample. Y.S., Q.H., J.L., Z.Z., and S.L. collected and processed the EM data. Y.S., Q.H., D.D., and Y.C. analyzed the structures. Y.S., Q.H., H.Y. and S.L. wrote the manuscript. All authors critically revised the manuscript.

## Funding

This work was supported in part from National Natural Science Foundation of China #32241031 and #32171195 (S.L.), Tsinghua University Spring Breeze Fund #2021Z99CFZ004 and Dushi Fund #2023Z11DSZ001 (S.L.), Fundamental Research Funds for the Central Universities #2022ZFJH003 (H.Y.), Zhejiang Plan for the Special Support for Top-notch Talents #2022R52029 (H.Y.), National Key Research and Development Program in China #2024YFC2309900 and #2021YFC2301204 (H.Y.).

### Date and materials availability

Electron microscopy maps have been deposited in the Electron Microscopy Data Bank under accession codes EMD-XXXXX, EMD-XXXXX, EMD-XXXXX, and EMD-XXXXX.

