## Supplementary Figures for "Cryo-ET of IgG bivalent binding on SARS-CoV-2 provides structural basis for antibody avidity"

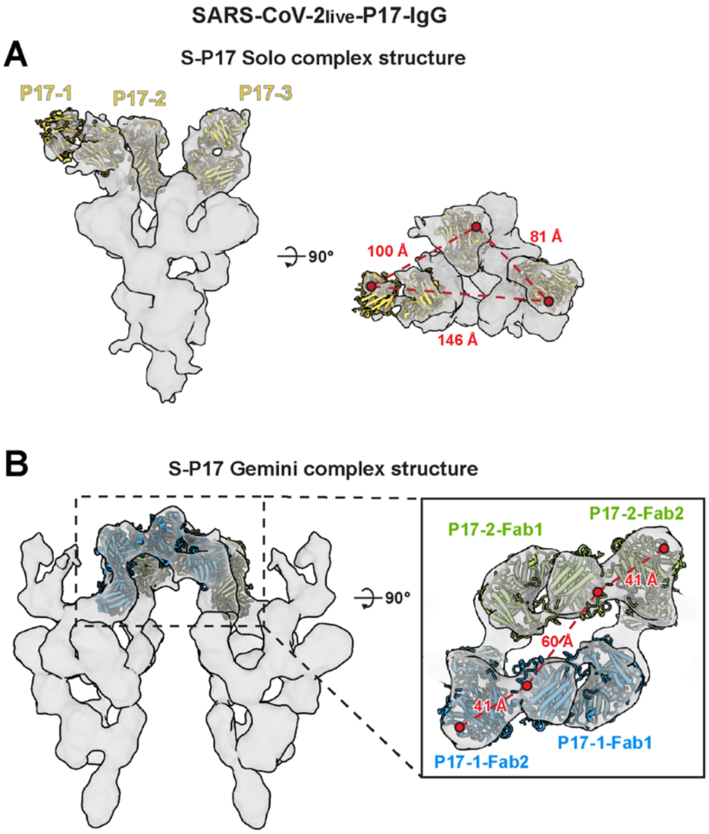


**Fig. S1. Spatial relationships of Fab fragments in the S-P17 Solo and Gemini complexes on SARS-CoV-2$_{live}$-P17-IgG.**

(**A**) Fab fragment models from PDB: 7K8O were fitted to the Fab densities of the S-P17 Solo complex. The measured distances between the C-terminal CH1 domains (residue 222 of the heavy chain) of adjacent Fab models were 81 Å, 100 Å, and 146 Å, respectively. (**B**) Four Fab models from PDB: 7K8O were fitted to the Fab densities bridging two adjacent S-trimers in the mode II(1) S-P17 Gemini complex. Two Fabs from one P17-IgG molecule (P17-1-Fab1 and P17-1-Fab2) were colored blue, while those from the second P17-IgG were colored light green. The measured distances between the CH1 domain C-termini were 41 Å for Fab1-Fab2 and 60 Å for Fab1-Fab1.

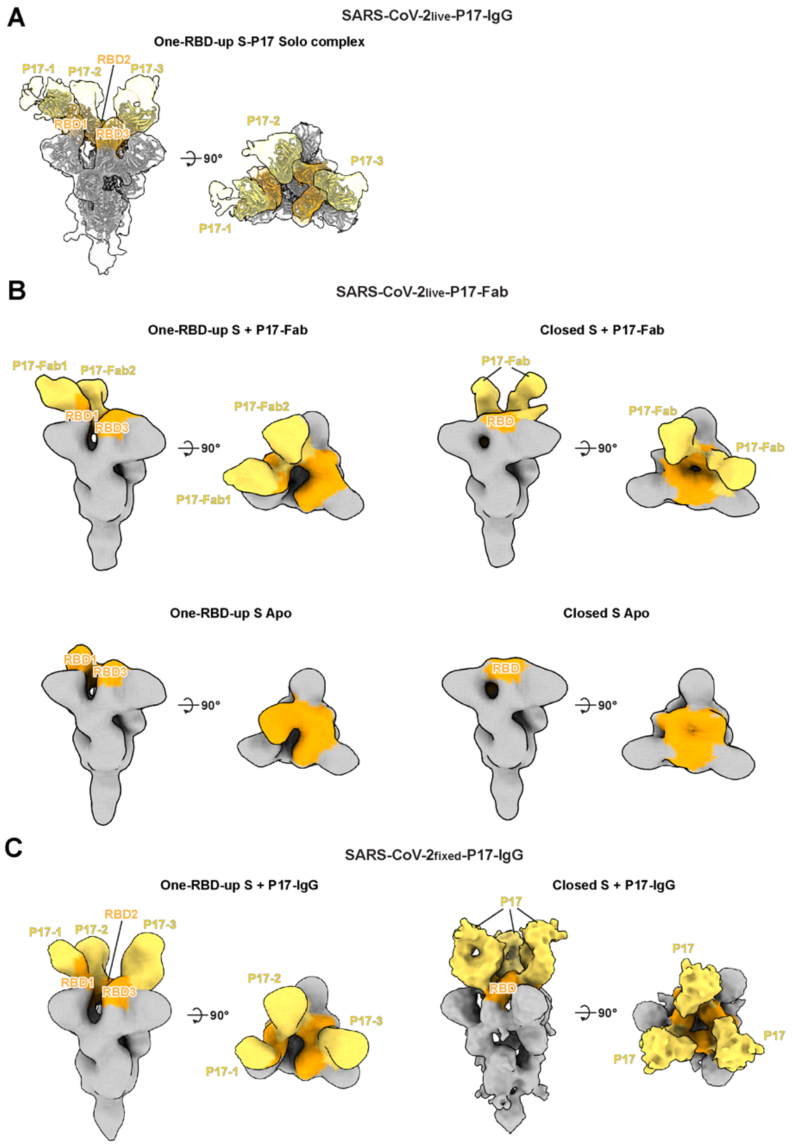


**Fig. S2. Structural analysis of S-P17 Solo complexes under varying conditions.**

(**A**) In the SARS-CoV-2$_{live}$-P17-IgG sample, all S-P17 Solo complexes adopt a one-RBD-up conformation, with each RBD bound by P17-IgG. A model of recombinant S engaging with P17-Fab (PDB: 7CWM) was fitted to the structure. (**B**) In the SARS-CoV-2$_{live}$-P17-Fab sample, four types of S-P17 Solo complex structures were identified: (1) one-RBD-up S with partial RBDs bound to P17-Fab, (2) closed S with partial RBDs bound to P17-Fab, (3) one-RBD-up S without P17-Fab binding, and (4) closed S without P17-Fab binding. (**C**) In the SARS-CoV-2$_{fixed}$-P17-IgG sample, S-P17 Solo complexes exist in both one-RBD-up and closed conformations, with all RBDs bound to P17-IgG. Densities corresponding to RBD and P17 were colored orange and yellow, respectively.

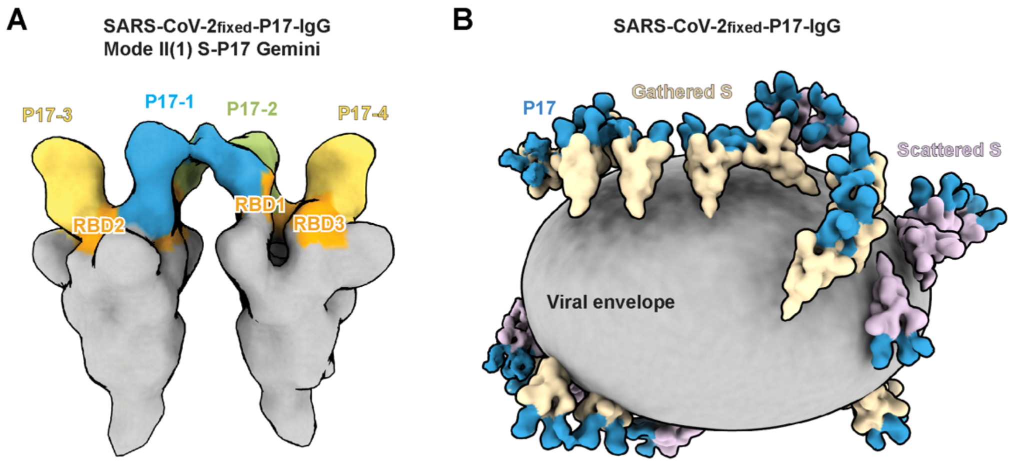


**Fig. S3. S-P17 Gemini complex structure and virion composite structure reconstructed from the SARS-CoV-2$_{fixed}$-P17-IgG sample.**

(**A**) A mode II(1) S-P17 Gemini complex structure determined from SARS-CoV-2$_{fixed}$-P17-IgG samples across all incubation durations. The structure shows two S-trimers (gray) with their RBDs (orange) bridged by two P17-IgG molecules (blue and green), similar to that of SARS-CoV-2$_{live}$-S309-IgG. (**B**) An exemplary SARS-CoV-2$_{fixed}$-P17-IgG virion was reconstructed by projecting all S-P17 Solo and mode II(1) S-P17 Gemini complexes determined from SARS-CoV-2$_{fixed}$-P17-IgG onto their refined coordinates on the viral envelope. The gathered S-trimers are colored wheat, while scattered S-trimers are colored thistle. P17 molecules (light blue) bridge the S-trimers, gathering them into patches on the viral surface. The viral envelope (gray) was reconstructed by manually segmenting membrane densities.

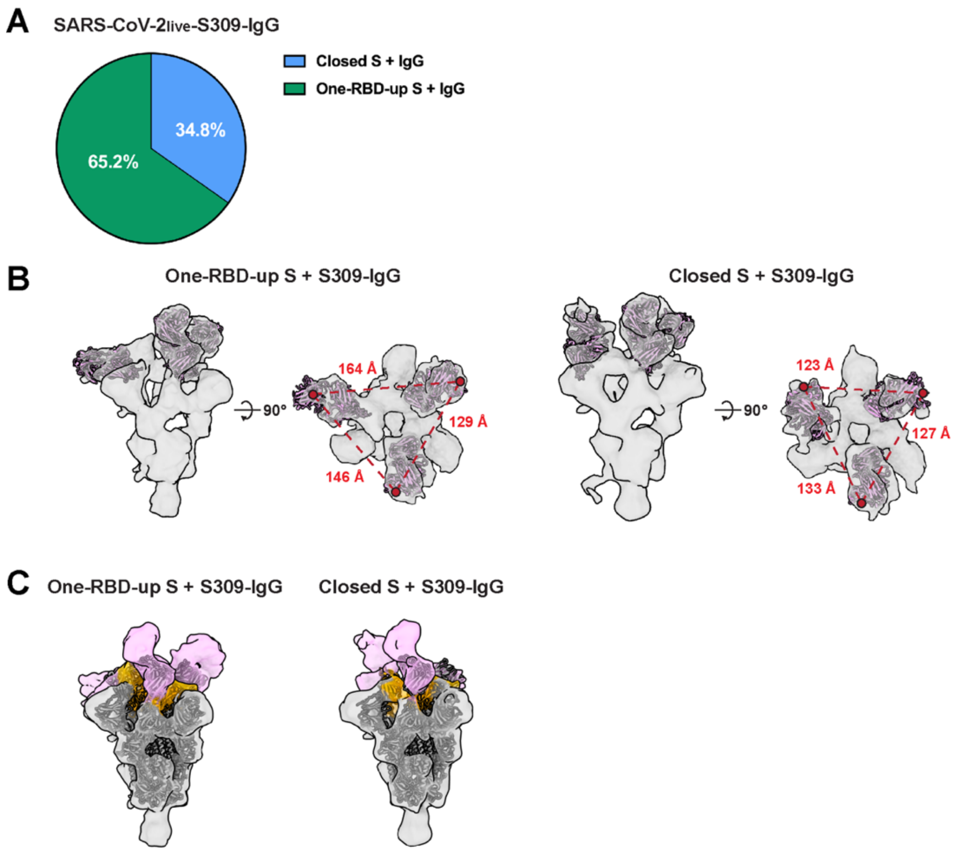


**Fig. S4. Structural characteristics of S-S309 Solo complexes in the SARS-CoV-$2_{live}$-S309-IgG sample.**

(**A**) In the SARS-CoV-$2_{live}$-S309-IgG sample, 65.2% of the prefusion S-trimers adopted a one-RBD-up conformation, while 34.8% adopted a closed conformation. (**B**) S309 Fab models from PDB: 6WS6 were fitted to the Fab densities of the S-S309 Solo complexes in both conformations. The measured distances between the C-terminal CH1 domains (residue 222 of the heavy chain) in the one-RBD-up conformation were 129 Å, 146 Å, and 164 Å. In the closed conformation, the distances were 123 Å, 133 Å, and 127 Å. (**C**) Models of recombinant S engaging with S309-Fab (PDB: 6WPT and 6WPS) were fitted to the S-S309 Solo complex structures. The densities of the RBD (orange) and S309-IgG (pink) in both the one-RBD-up and closed conformations align well with the recombinant structures.

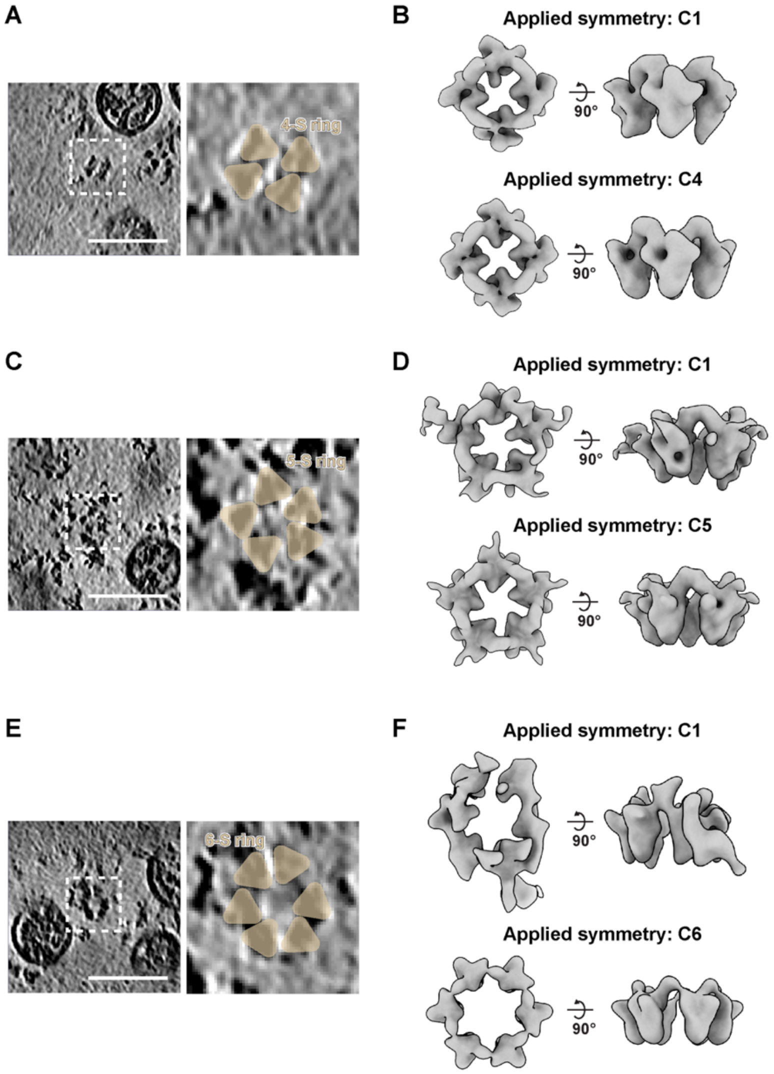


**Fig. S5. STA results of the circular S-S309 assemblies in SARS-CoV-2$_{live}$-S309-IgG.**
(**A**) An exemplary tomogram slice showing a tetramer-of-trimers structure (white box), where each S density is labeled as a wheat-colored triangle. (**B**) STA of 69 tetramers-of-trimers assemblies was performed, applying either C1 or C4 symmetry. (**C**) An exemplary tomogram slice showing a pentamer-of-trimers structure (white box). (**D**) STA of 53 pentamers-of-trimers applying either C1 or C5 symmetry. (**E**) An exemplary tomogram slice showing a hexamer-of-trimers structure (white box). (**F**) STA of 17 hexamer-of-trimers applying either C1 or C6 symmetry. Tomogram slice thickness: 5 nm; scale bar: 100 nm.

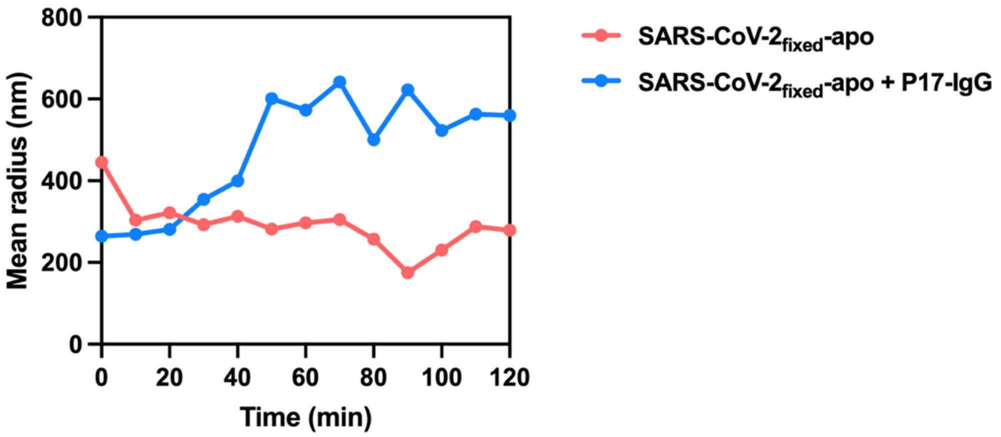


**Fig. S6. Characterization of virion aggregation induced by P17-IgG via dynamic light scattering (DLS).**

The mean hydrodynamic radius of the SARS-CoV-$2_{fixed}$-apo sample increased over time following the introduction of P17-IgG, indicating progressive virion aggregation. In contrast, the hydrodynamic radius of the SARS-CoV-$2_{fixed}$-apo samples without P17-IgG introduction remained relatively stable.

### Table S1. Cryo-ET data acquisition and reconstruction statistics of P17.

| Data acquisition | | | | | | | | | |
|---|---|---|---|---|---|---|---|---|---|
| Microscope | Titan Krios | | | | | | | | |
| Magnification | 64,000 | | | | | | | | |
| Voltage (kV) | 300 | | | | | | | | |
| Detector | Gatan K3 | | | | | | | | |
| Energy filter (eV) | 20 | | | | | | | | |
| Pixel size (Å) | 0.68 (super-resolution) | | | | | | | | |
| Tilt schemes | Dose-symmetric/Bidirectional scheme | | | | | | | | |
| Exposure (e$^-$/Å$^2$) | 131.2 | | | | | | | | |
| Defocus range (μm) | -2.0 ~ -4.0 | | | | | | | | |
| Software | SerialEM | | | | | | | | |
| **Reconstruction** | | | | | | | | | |
| Software | Dynamo 1.1.333 | | | | | | | | |
| Samples | SARS-CoV-2$_{live}$-P17-IgG | | SARS-CoV-2$_{fixed}$-P17-IgG | | | SARS-CoV-2$_{live}$-P17-Fab | | | |
| No. of tomograms | 276 | | 60 | | | 54 | | | |
| No. of virions | 3,258 | | 885 | | | 599 | | | |
| Structure datasets | S-P17 Solo (One-RBD-up) | Mode II(1) S-P17 Gemini | S-P17 Solo (Closed) | S-P17 Solo (One-RBD-up) | Mode II(1) S-P17 Gemini | S-P17-Fab (Closed) | S-P17-Fab (One-RBD-up) | S-apo (Closed) | S-apo (One-RBD-up) |
| Final no. of particles | 9,246 | 9,835 | 4,628 | 4,884 | 818 | 1,822 | 1,825 | 6,086 | 2,477 |
| Symmetry imposed | C1 | C2 | C3 | C1 | C2 | C1 | C1 | C1 | C1 |
| Final Resolution (Å) | 11.4 | 12.4 | 11.4 | N/A | N/A | N/A | N/A | N/A | N/A |
| Gold-standard | yes | yes | yes | no | no | no | no | no | no |
| FSC threshold | 0.143 | 0.143 | 0.143 | N/A | N/A | N/A | N/A | N/A | N/A |
| Final pixelsize (Å) | 2.72 | 2.72 | 2.72 | 5.44 | 5.44 | 5.44 | 5.44 | 5.44 | 5.44 |

**Table S2. Cryo-ET data acquisition and reconstruction statistics of S309.**

| **Data acquisition** | | | | | | | | | | |
|---|---|---|---|---|---|---|---|---|---|---|
| Microscope | Titan Krios | | | | | | | | | |
| Magnification | 64,000 | | | | | | | | | |
| Voltage (kV) | 300 | | | | | | | | | |
| Detector | Gatan K3 | | | | | | | | | |
| Energy filter (eV) | 20 | | | | | | | | | |
| Pixel size (Å) | 0.68 (super-resolution) | | | | | | | | | |
| Tilt schemes | Dose-symmetric | | | | | | | | | |
| Exposure ($e^-$/Å$^2$) | 131.2 | | | | | | | | | |
| Defocus range (µm) | -2.5 ~ -3.5 | | | | | | | | | |
| Software | SerialEM | | | | | | | | | |
| **Reconstruction** | | | | | | | | | | |
| Software | Dynamo 1.1.333 | | | | | | | | | |
| Samples | SARS-CoV-2$_{live}$-S309-IgG | | | | | | | | | |
| No. of tomograms | 33 | | | | | | | | | |
| No. of virions | 492 | | | | | | | | | |
| Structure datasets | S-S309 Solo (Closed) | S-S309 Solo (One-RBD-up) | Mode (1) S-S309 Gemini | Mode (2) S-S309 Gemini | Mode (3) S-S309 Gemini | Mode (4) S-S309 Gemini | Mode (5) S-S309 Gemini | S-S309 tetramer | S-S309 pentamer | S-S309 hexamer |
| Final no. of particles | 2,153 | 4,237 | 1,045 | 1,099 | 1,509 | 572 | 433 | 69 | 53 | 17 |
| Symmetry imposed | C1 | C1 | C2 | C1 | C1 | C1 | C1 | C4 | C5 | C6 |
| Final Resolution (Å) | 16.0 | 14.5 | 14.4 | N/A | N/A | N/A | N/A | N/A | N/A | N/A |
| Gold-standard | yes | yes | yes | no | no | no | no | no | no | no |
| FSC threshold | 0.143 | 0.143 | 0.143 | N/A | N/A | N/A | N/A | N/A | N/A | N/A |
| Final pixelsize (Å) | 2.72 | 2.72 | 2.72 | 5.44 | 5.44 | 5.44 | 5.44 | 5.44 | 5.44 | 5.44 |
